# Human Action Planning and Mental Rotation in a Tetris-Like-Game

**DOI:** 10.1101/2025.09.28.679040

**Authors:** Aisha Aamir, Minija Tamosiunaite, Florentin Wörgötter

**Affiliations:** Göttingen University, Institute for Physics 3 – Biophysics, and Bernstein Center for Computational Neuroscience, Göttingen, Germany; Vytautas Magnus University, Department of Informatics, Vileikos 8, 44158 Kaunas, Lithuania

## Abstract

The mechanisms behind human action planning and mental object manipulation are still not well understood. These core cognitive abilities are essential not only for interaction with physical spaces, e.g., for assembling objects, but also for effective problem-solving in the digital world. Here we ask, which strategies humans employ when assessing whether or not an object will fit into a cavity. To this end, objects and cavities were presented with different orientations on a computer screen and we measured errors, reaction-times and gaze patterns, where the latter can point to different problem-solving strategies. On the one hand our findings confirm that simpler configurations are solved faster and more efficiently than more complex ones. On the other hand, by analyzing about 80,000 gazes, we observed that participants used three different strategies. In many instances, the investigated task — featuring relatively large objects — could be completed using only peripheral vision (37%). In a larger number of cases quite “specific” gaze patterns were observed, primarily focusing on the Gestalt of a concave corner (46%). Less frequently, but still notably often, participants employed a strategy of fixating near to object or cavity (17%), potentially minimizing the length of the required saccadic eye movements while relying on perifoveal/peripheral vision. Ultimately, these findings highlight the crucial roles of proximity, spatial orientation, and visual cues in object recognition tasks, suggesting that the perceptual strategies used depend on distinct aspects of the object configurations.

## 1. Introduction

In the fast-evolving field of cognitive science, the processes underlying human action planning and mental object manipulation remain a compelling challenge. These cognitive functions are not only fundamental for navigating our spaces and assembling objects but are also critical for problem-solving in dynamic digital environments (Xue et al., 2017). An exemplary paradigm to study these functions is found in the Tetris game, which demands rapid decision-making, anticipatory planning, and spatial manipulation of objects. When playing, falling blocks with different shapes and orientations on a computer screen have to be sorted so as to form a compact ground plane. As blocks fall at varying speeds, players must mentally rotate and position them while simultaneously coordinating a sequence of motor actions and visual evaluations (Gentile and Lieto, 2022).

The significance of mental rotation—the ability to manipulate mental representations of objects—has been highlighted throughout decades of spatial cognition research. Classic experiments indicate that the time required to perform a mental rotation increases linearly with the angle of rotation. This finding provides insights into the cognitive load and processing strategies that underpin task performance (Ågren et al., 2023). However, most prior research has focused on static or sequential presentations of stimuli, which leaves a considerable gap in understanding how these mechanisms function in the context of a continuously evolving environment.

Tetris provides a rich experimental platform for studying action planning and mental rotation, both of which are key cognitive processes involved in dynamic problem-solving. Players must quickly assess falling tetrominoes^1^, anticipate their placement within the existing structure, and decide how to rotate them for optimal fit. This requires efficient spatial reasoning and decision-making under time constraints.

In this regard many studies have been conducted; for instance, Gentile and Lieto (2022) present an innovative ACT-R computational model to explore how mental rotation processes are activated during Tetris gameplay. The authors demonstrate that mental rotation is not continuously active, but rather selectively engaged depending on the game dynamics and task demands. Their model integrates insights from spatial cognition and cognitive architecture research, revealing that effective decision-making in Tetris hinges on the timely activation of mental rotation processes.

Using a different setup, Thomas and Lleras (2007) explore the implicit link between eye movements and spatial cognition, demonstrating that gaze trajectories can influence problem-solving efficiency. Their study reveals that participants who unconsciously moved their eyes in patterns aligned with a problem’s solution were more successful in solving it, suggesting spatial compatibility between visual exploration and cognitive processing. The findings support the idea that eye movements are not merely passive reflections of thought but actively shape reasoning strategies.

Kirsh and Maglio (1994) introduce the distinction between epistemic actions and pragmatic actions in cognitive science, using again Tetris as a case study to illustrate how physical actions can enhance cognitive processing. They argue that epistemic actions — such as rotating a Tetris block before deciding its placement — serve to simplify mental computation. In contrast, pragmatic actions are performed to physically progress toward a solution. Their findings challenge traditional models that assume all actions are goal-directed, highlighting how external manipulations can reduce cognitive load and improve problem-solving efficiency. Hsing and Bairaktarova (2022) investigate how eye-tracking can uncover cognitive processes and problem-solving strategies used by engineering students during spatial rotation and mental cutting tasks. Their study categorizes gaze behavior into encoding, transformation, and confirmation processes, revealing that encoding fixations are strongly correlated with accuracy and strategy selection. Findings suggest that mental rotation tasks require more piecemeal strategies, while mental cutting tasks favor holistic approaches. Ågren et al. (2023) investigate the neural mechanisms underlying Tetris gameplay, focusing on its visuospatial processing demands. Using functional magnetic resonance imaging (fMRI), the study identifies activation in the ventral and dorsal visual streams, particularly in the inferior and mid-temporal gyrus, when players engage in mental rotation tasks during gameplay. Their findings suggest that Tetris recruits brain regions associated with spatial reasoning and working memory; supporting its use in psychological interventions targeting intrusive mental imagery and cravings. Lindstedt and Gray (2015) introduce Meta-T, a modified Tetris paradigm designed for cognitive skills research, offering precise experimental control over decision-making, perceptual-motor interactions, and time-stressed problem-solving. Unlike traditional gaming studies, Meta-T provides detailed logging of user actions, enabling researchers to analyze real-time cognitive adaptations. Xue et al. (2017) investigate the cognitive processes underlying mental rotation using eye-tracking data to analyze how individuals engage with spatial transformations. Their study employs a discriminative Hidden Markov model (dHMM) to segment mental rotation into different distinct processing states. The intersection of Tetris gameplay, eye tracking, and computer vision represents a promising multidisciplinary approach to understanding and facilitating action planning. Krol, et al. (2017) further illustrated that when manual inputs are replaced with gaze-based control—even in a complex dynamic environment like Tetris—the resulting patterns reveal important markers of cognitive load and strategic planning. These insights not only deepen our theoretical understanding of visuospatial processing but also point toward practical applications in the design of adaptive human–computer interfaces.

In this study, we investigate how cognitive skills, particularly eye movements relate to action as well as planning processes in humans. Our research uses a much-reduced version of the Tetris game, where one tetromino – called “Part” in the following – needs to be assesses whether it would fit (or not) into a cavity – called “Hole” – at the bottom of the display (see Figure 1). As participants ‘play’, their eye movements are tracked to identify patterns and correlations between gaze behaviors and decision-making outcomes. This approach enables us to quantify the cognitive load involved in the task, providing insights into how visual attention is allocated during gameplay. Based on this paradigm, we consider the following hypotheses:

**Figure 1.**
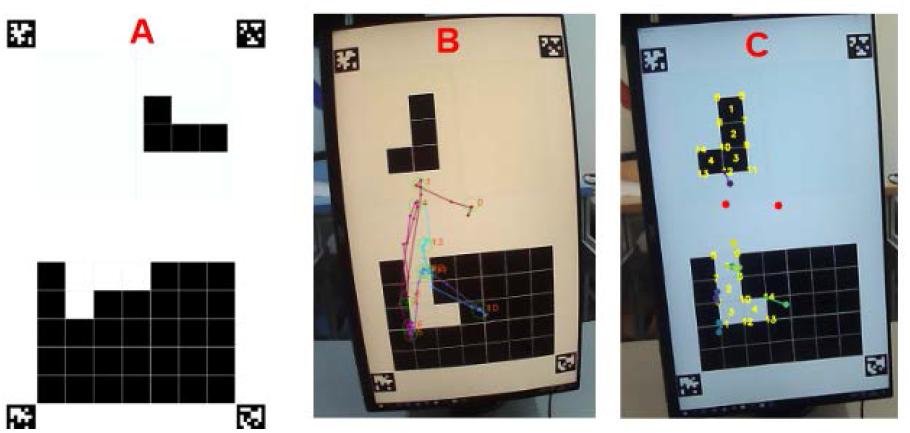
Screen setup. (A) Representation of a game situation as shown on the screen. (B) Example Part and Hole situation with eye-movement tracks and fixation point sequence (numbers) indicated on the still distorted image of the screen. Only long-enough fixations (> 300 ms) are considered (colored circles). (C) Same results but now showing characteristic markers at Part and Hole and fixation points (dots), where lines annotate the closest marker to a given fixation point. Red fixations are out of range for Part-Hole analysis and are treated separately. This panel shows the picture of the screen after distortion correction.

1. People will recognize a fit between Part and Hole faster and with less errors as compared to a non-fit, where simple configurations will lead to shorter reaction times than more complex ones. For example, the equivalent of the regular, upright letter L and its mirror image should be more easily correctly assigned (as “L” is a common entity for adult, literal humans).
2. Concerning gaze patterns, specific regions of the display and/or Gestalt-like configurations of Part and/or Hole might matter.

## 2. Methods

### 2.1 Participants

The current study involved 40 participants, consisting of 25 males and 15 females, with an average age of 24.4 years and a standard deviation of 2.7 years. Every participant had regular vision or was wearing contact lenses. Participation was voluntary, and prior to engaging in the experiment, each individual was provided with a clear and concise introduction to the study’s objectives. Informed consent was obtained from all participants and kept on record, ensuring that they were fully aware of their rights and the nature of the experiment. The experiment is not harmful and no sensitive data had been recorded, experimental data has been treated anonymously and only the instructions explained below had been given to the participants. Participants were allowed to stop the experiment in case of discomfort or fatigue. The experiment was performed in accordance with the ethical standards laid down by the 1964 Declaration of Helsinki. We followed the relevant guidelines of the DPG (Document: 28.09.2004 DPG: “Revision der auf die Forschung bezogenen ethischen Richtlinien”) and these experimental procedures had been assessed by the Ethics Committee of the University of Göttingen, Psychology. Approval has been obtained under registration number 256.

### 2.2 Tools and Software

In our experiment, we used eye tracker glasses from Pupil Labs, with software version 3.5.1. This configuration operates at a camera frame rate of 60 Hz, allowing for precise data collection and analysis of gaze patterns.

### 2.3 GUI Configuration and Puzzle Pieces

We developed the game interface using Python PyQt/Qml packages, prioritizing simplicity and neutral colors to ensure the design does not interfere with the goals of the experiment. The game’s GUI features a base block in black with a “Hole” with a white canvas above where different incoming pieces, the “Parts”, are presented (Figure 1). Each round introduces a random combination of Hole and Part. The design orientation is in portrait-format on a vertically oriented computer monitor to ensure optimal spatial resolution with sufficiently large configurations for accurate eye tracking. Participants were asked to indicate whether the Part would fit into the Hole. To indicate this, participants used the “M” key to indicate fit, the “Z” key for not fit and the Spacebar to start the game. The next configuration was then shown 600 ms later.

The game features 12 puzzle pieces named L and LR (‘R’ for reversed, see Figure 2) as well as Z and ZR. The numerical identifiers denote orientation; for example, the L puzzle has four rotations labeled L_0, L_90, L_180, and L_270. All potential rotations with their corresponding IDs are presented in Figure 2.

**Figure 2.**
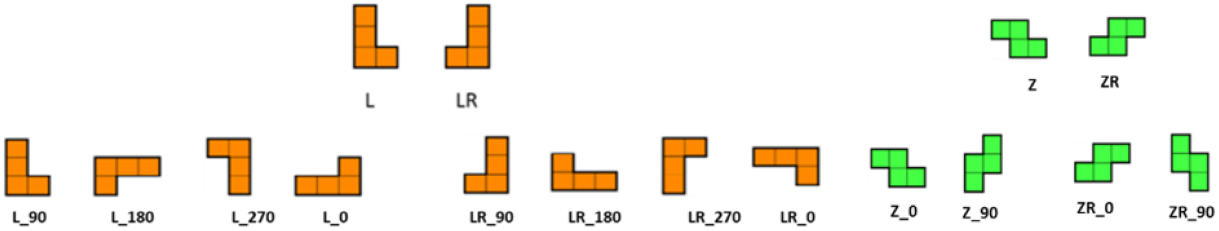
L,LR, Z and ZR puzzle pieces and their possible allowed rotations and IDs used in the current Tetris-like game experiment.

In an experiment, every participant sees multiple rounds featuring various Part and Hole combinations, as illustrated in Figure 1, where Parts and Holes can be at different locations on the screen. After the user has pressed the “Fit” or “Not Fit” button, the next round is started with a new combination of

Part and Hole. For each round, reaction times and classification outcomes are recorded together with all eye-movements.

### 2.4 Gaze Matching

This process begins with the transformation of raw eye-tracking data from the video coordinate system to an undistorted image space, ultimately converting it to screen pixels. Once the eye-tracking data was converted, we aligned these readings with the designated Part and Hole locations identified by computer vision methods (see Appendix). Notably, each fixation leads to multiple recorded sampling data points and by calculating the median x and y positions, we were able to extract from this the most reliable fixation location. The median was favored for its robustness against outliers, providing a clearer representation of where fixation was located. Figure 1b shows an example of this analysis projected back onto the (still distorted) picture of the screen.

Subsequently, we divided the screen into three regions, bottom, middle, and top and calculated the Euclidean distance between each fixation and the center and border positions of the Parts and Holes, as indicated by yellow markers in Figure 1c. To ensure accuracy, we applied a minimum distance threshold equal to the grid spacing size, considering only the nearest matches. This filtering process effectively captures the most relevant matches, clustering fixations at or around the different marker points.

### 2.5 Experimental Procedures

Before an experiment commenced, each participant calibrated the eye tracker camera mounted on the glasses. This calibration occurred on the vertically oriented computer screen, where calibration markers (small circles) were displayed at the center and all four corners of the canvas. Participants were instructed to maintain a still head position while looking at the circles with their eyes. This process ensures highly accurate calibration for measuring eye movements throughout the experiment. To further enhance accuracy, a calibration circle, onto which participant had to fixate, was introduced after every round, ensuring consistent starting points for fixations across the experiment.

We conducted two studies. The first utilized the full set of 12 Parts and focused on analyzing reaction times and classification errors with 10 participants, without recording eye movements. The duration of these experiments was approx. 60 min and this proved too long for participants making it difficult to achieve precise gaze recordings. Therefore, to shorten the experiment duration, only L-Types were selected for the second study. This choice was informed by the results of the first study, which indicated that L-Types exhibited more complex behavior compared to Z-Types.

In the first study, we used 104 combinations of L, LR, Z, and ZR pieces. Within the same L or Z class, all Part-Hole combinations were included, whereas for control conditions (L vs. Z and Z vs. L), only two Holes from the opposite class were used. The 104 combinations were presented to participants in a pseudo-random order, with each combination repeated four times, resulting in a total of 416 rounds.

For the second study, we reduced the number of potential combinations to 64, leading to a total of 256 rounds and a shorter experiment duration of less than 30 minutes. Fewer intermittent calibrations were needed, further streamlining the process. For this we recorded data from 30 participants.

To ensure consistency across participants, the sequence of combinations was stored in a JSON file. Participant data—including Part- and Hole-IDs, reaction times, and classification results—was recorded in an Excel sheet. Additionally, for the second study, all eye movements were tracked (see Appendix for details).

## 3. Results

We present the results in two parts following the two different experimental designs.

### 3.1 Experiment 1: Reaction times and errors

This study measured reaction times and error rates for 10 participants solving puzzle combinations L, LR, Z, and ZR. The data is annotated in tuples: (Part, Hole), and we consider all possible pairings: (L, L), (L, LR), (LR, L), (LR, LR), (Z, Z), (ZR, Z), (Z, ZR), and (ZR, ZR). However, we aggregate all data from the following groups: (L, any Z), (LR, any Z), (Z, any L), and (ZR, any L). The distinction between L-type and Z-type configurations is pronounced enough to serve as “controls” in this study. As illustrated in Figure 3, reaction times for these four control-cases are consistently lower and exhibit minimal dispersion compared to the others.

**Figure 3.**
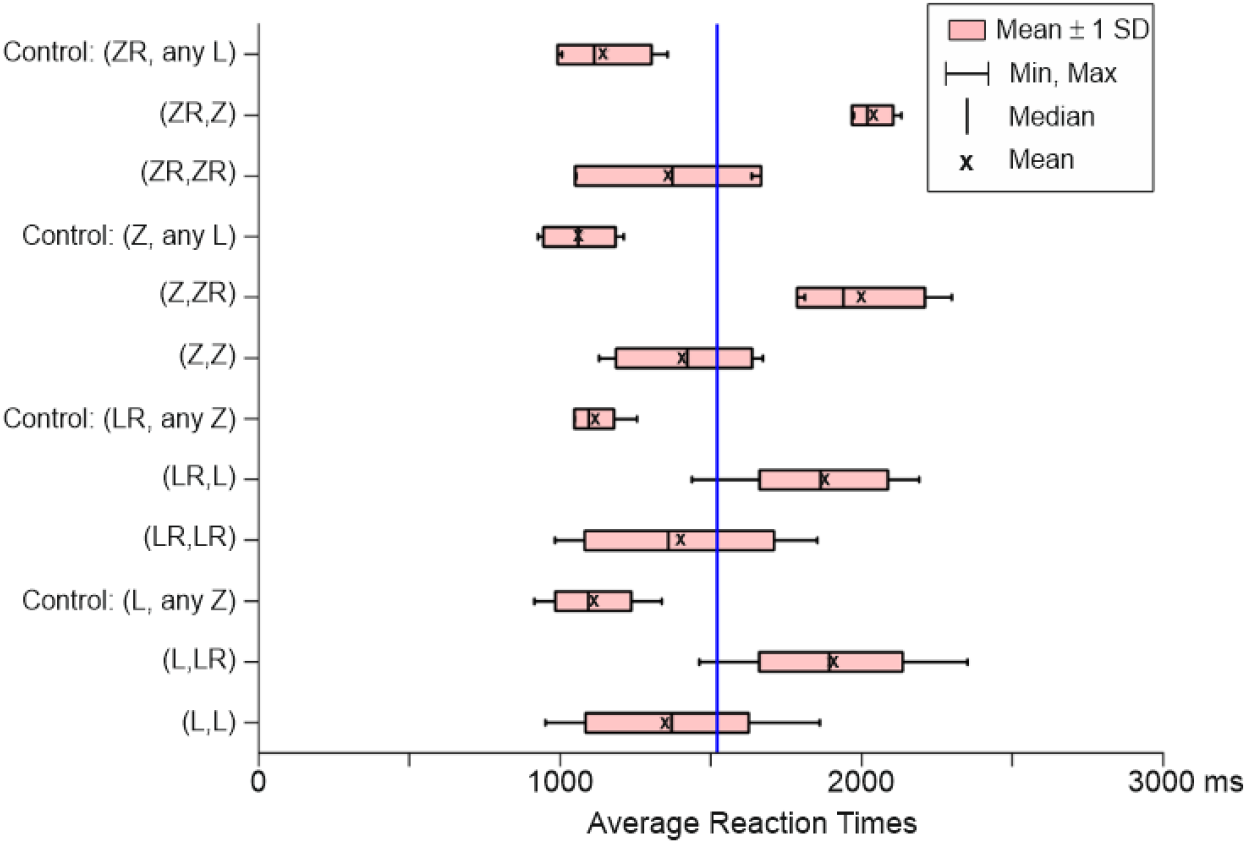
Average reaction time (ms) for 10 participants in Study 1. The game was designed to include various puzzle combinations. The blue line represents the grand average across all experiments

Another group of four cases is that of equally-paired combinations (L,L), (LR,LR), (Z,Z), and (ZR,ZR). They lead to reaction times smaller than those of the last group – the unequally-paired combinations (ZR,Z), (Z,ZR), (L,LR), and (LR,L). Interestingly, data dispersion is higher for the equally-paired than for the unequally-paired group.

Table 1 presents the mean and standard deviations for equally-paired versus unequally-paired combinations, providing a clearer visualization of the results discussed earlier. Generally, the values on the right side of the table—representing the equally-paired cases—are lower than those on the left. This difference is statistically significant (p < 0.001) across all cases. The average reaction time for all control cases is approximately 1100 ms, with relatively small standard deviations. These averages are, therefore, significantly lower (p < 0.001) than those observed in all other cases in Table 1.

**Table 1.** Mean and standard deviations for equally-paired versus unequally-paired combinations. Top: Z-types, bottom: L-types.

| Mean | SD | Mean | SD |
| --- | --- | --- | --- |
| **Control (Z, any L)** |  | **Control (ZR, any L)** |  |
| 1137 | 161 | 1073 | 65 |
| **(ZR,Z)** |  | **(ZR,ZR)** |  |
| 2036 | 59 | 1358 | 268 |
| **(Z,ZR)** |  | **(Z,Z)** |  |
| 1975 | 148 | 1410 | 195 |
| **Control (L, any Z)** |  | **Control (LR, any Z)** |  |
| 1161 | 94 | 1060 | 61 |
| **(LR,L)** |  | **(LR,LR)** |  |
| 1874 | 206 | 1396 | 304 |
| **(L,LR)** |  | **(L,L)** |  |
| 1898 | 230 | 1354 | 262 |

### 3.2 Classification Error Rates

Figure 4 presents a visual representation of error rates for each puzzle configuration. Blue and red indicate cases where participants made no errors in determining whether a Part fits the Hole. For all other cells, the classification error rate is displayed both numerically as a binned percentage and visually through a color gradient. Notably, across all experiments, only a single error was recorded for the control cases, and these squares are therefore excluded from the figure.

**Figure 4.**
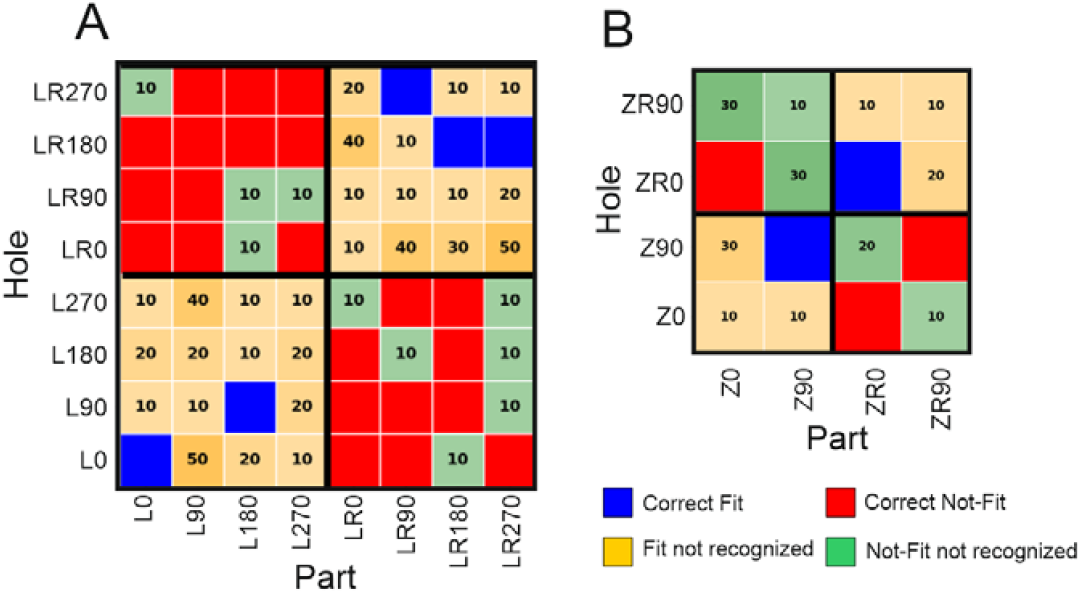
Classification results with binned error rates across participants. Binning in %, e.g., [0,10], [10,20], etc. (A) L-types. (B) Z-types.

In an error-free recognition scenario, the map would display the two equally-paired quadrants (lower left and upper right) in blue, indicating correct fits, while the remaining quadrants would be red, signifying correct non-fits. For L-types, correct non-fits were more frequent than correct fits, resulting in more red than blue. As explained earlier, this correlates with longer reaction times in these instances. Errors along the diagonal of equal-orientation pairs (bottom-left to top-right) were slightly lower than elsewhere in these quadrants, a pattern observed for both L-types and Z-types. However, the error distribution for Z-types was more balanced than for L-types, with errors occurring at similar frequencies across all quadrants.

Additionally, we examined for the L-types, for which more combinations exist than for the Z-types, whether there is a correlation between reaction times and correct or incorrect decisions. This data is not presented, as no significant effects were found. We observed only a non-significant trend of shorter reaction times for correct decisions compared to incorrect ones and that the reaction times’ standard deviations for incorrect decisions are larger than those for correct ones.

### 3.3 Experiment 2: Center of mass vs. specific detail gazes

In the next step, we analyzed participants’ detailed gaze patterns using data from the second study, which exclusively presented L-Type stimuli. In total, we examined N_tot_ = 81,961 gazes. Our primary interest was the specificity of gaze patterns i.e., how often participants focused on particular Part/Hole features compared to making more general fixations? To investigate this, we conducted a two-step data split. First, we identified unspecific gazes directed toward the center of the canvas and separated them from gazes more directly targeting the Part/Hole. In the second step, we examined the distribution of these more specific gazes to determine precisely where they were directed.

Split 1: We recorded a total of N_tot_ = 81,961 fixations. To classify the gazes, we divided the canvas into three equal regions: top, middle, and bottom, dividing the distance between Part and Hole into 3 times 1/3^rd^ (Figure 5b). For example, gaze 0 in Figure 5b falls within the middle region, while all others can be attributed to Part or Hole. See also Figure 1 where two red fixations are marked as falling in the middle, according to this definition.

**Figure 5.**
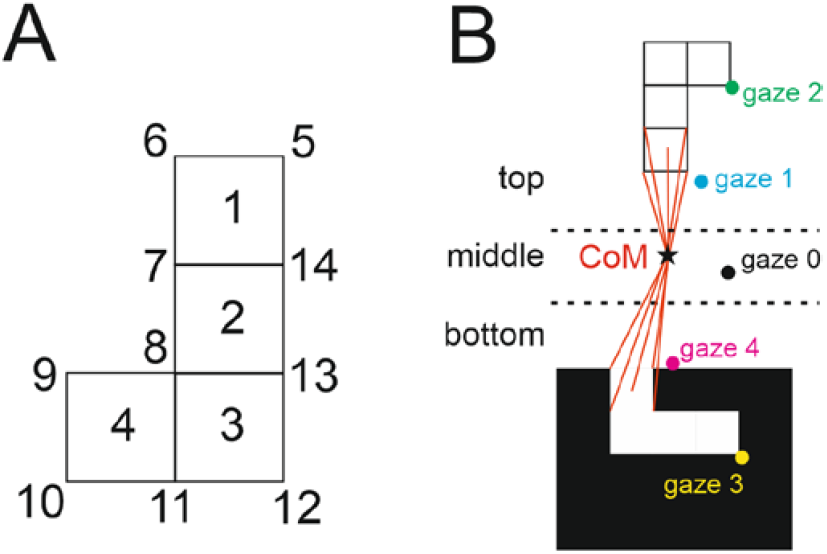
Marking of specific features of a hypothetical Part/Hole configuration with a) showing the 14 marker points and b) Divisions of the canvas and center of mass (CoM).

Overall, 37% of all gazes landed in the middle region. In the next step, we excluded these middle-region gazes (N_middle_ = 30,636) and analyzed only the gazes clearly targeting Part or Hole (i.e., non-middle) where we had N_not-middle_ = 51,325 gazes.

Split 2: This split is motivated by the question whether participants might sometimes judge a scene by looking “just close enough” to the Part/Hole without focusing on specific features such as line segments or corners. Since larger saccades generally require more effort than smaller ones, it follows that these “close-enough to Part/Hole” gazes would most likely occur near the bottom of the Part or near the top of the Hole. This positioning allows alternating gaze patterns between the Part and Hole to remain relatively effortless.

To establish a quantitative method, we assigned 14 marker points to both, Parts and Holes, as illustrated in Figure 5a. Additionally, to define a reference point for analyzing gaze patterns, we defined the center of mass (CoM), calculated by treating the blocks of Part and the empty blocks of Hole equally, like entities with mass, and determining the CoM then as usual. The CoM for the configuration in Figure 5b is schematically depicted as a black star.

The CoM can then be utilized to analyze different gaze patterns, specifically examining how many gazes are directed *at markers that are close to the CoM* versus those that are farther away. In Figure 5b, red vectors highlight the five closest markers to the CoM for both, Part and Hole. For each scenario, we calculated all 2 × 14 vectors (some are shown in red in Figure 5B), allowing us to determine, based on the specific situation, which markers are near the CoM and which are not.

In the given example, the ordering of the five closest markers, from nearest to farthest, is as follows: for the Part: 5, 6, 1, 7, 14; and for the Hole: 9, 10, 4, 8, 11 (see Figure 5a for marker numbers). As shown in Figure 5b, gaze 1 is close to marker 6 on the Part, and gaze 4 is near marker 9 on the Hole, whereas gazes 2 and 3 are positioned farther away.

The question remains: how can we quantify what constitutes “looking just close to” the Part/Hole versus “far”? To address this, we utilize this representation to compute a rank-order histogram, capturing how frequently participants fixate on the closest marker, the second closest, the third closest, and so on relative to the CoM. Figure 6 presents the average results for all 30 participants across both Parts and Holes.

**Figure 6.**
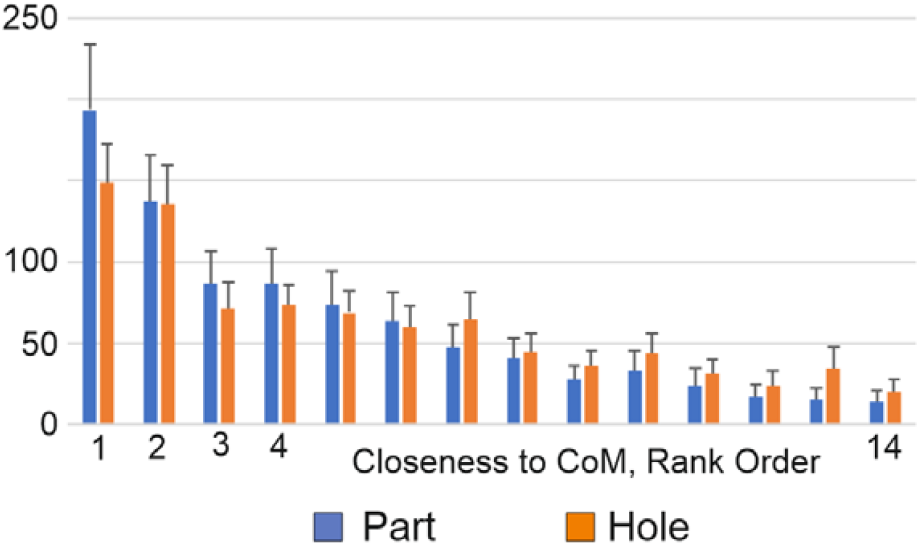
Rank-order histogram capturing how frequently participants fixate on the closest marker, the second closest, the third closest, a3n0d0so on relative to the CoM, average over 30 participants and confiden intervals of the mean are provided.

This figure shows that the markers of rank 1 and 2, which are the ones closest-to and 2^nd^-closest-to the CoM, are looked at rather often, after which there is a clear drop in the histogram. This drop is significant as shows by the fact that the score of rank 2 is above the confidence intervals of rank 3 (and higher). For ranks 3 and higher the histogram only gradually decreases.

This suggest that Participants do indeed often look “just close-enough” at Part or Hole, regardless of the actual Part/Hole configuration, corresponding to gazes covered in Figure 6 by rank 1 and 2. Hence we divided the data counting the number of gazes of rank 1 plus rank 2, which represent those that are “Close-to” Part or Hole, as compared to the rest, which we call “Specific” gazes (the reason for this naming convention will become clear later).

Table 2 shows this data split, where 25,704 gazes are in the top region (See Figure 5 b) and a similar number, 25,621, are in the bottom region of the canvas. Chance “Close-to” looks (ranks 1 and 2) would lead to an average of 1/7th of all cases (=7,332). However, Participants look “Close-to” in more than 1/4th of all cases and they look “Close-to” Parts significantly more often than “Close-to” Holes, X_2_(1, N=51,325) = 1,770.12, p<0.0001.

**Table 2.** Gaze counts for different locations at the canvas and percent values for different data splits.

|  | Part | Hole | Sum | % | % | % |
| --- | --- | --- | --- | --- | --- | --- |
| Not Middle and Close-to | 8939 | 4706 | $N_{\text{Close-to}}=13645$ | | 26 | 17 |
| Not Middle and Specific | 16765 | 20915 | $N_{\text{Specific}}=37680$ | | 74 | 46 |
| Sum Not Middle | 25704 | 25621 | $N_{\text{not-middle}}=51325$ | 63 | 100 | |
| Middle | | | $N_{\text{middle}}=30636$ | 37 | | 37 |
| All=Middle + Not-middle | | | $N_{\text{total}} = 81961$ | 100 | | 100 |

Hence, there is a large number of gazes which are not directed specifically to any location of Part or Hole. These are N_middle_=30636 and N_Close-to_=13645 (=54% of the overall total, N_total_=81961) with a remaining N_Specific_=37680, where the latter still equals 46% of the overall total.

Figure 7 shows how often do individual participants look just “Close-to” versus “Specific” at Parts and Holes? For example, the rightmost two bars represent the fraction of participants who had looked “Specific” at Part (blue) or Hole (orange) with more than 80% of their gazes. The blue histogram shows that only one person performed more than 80% “Specific” gazes at Parts. All others made less and, accordingly, those were more often able to use just a “Close-to” look to judge a situation. This is different for Holes, where 19 Participants performed more than 80% “Specific” gazes. The whole histogram orange is shifted to the right relative to the blue one and, overall, more participants needed to look more-often “Specific” at the Hole to assess its orientation than at the Part. Furthermore, we observe that the leftmost bin remains empty. Hence, all Participants performed at least some “Specific” gazes at Part as well as Hole.

**Figure 7.**
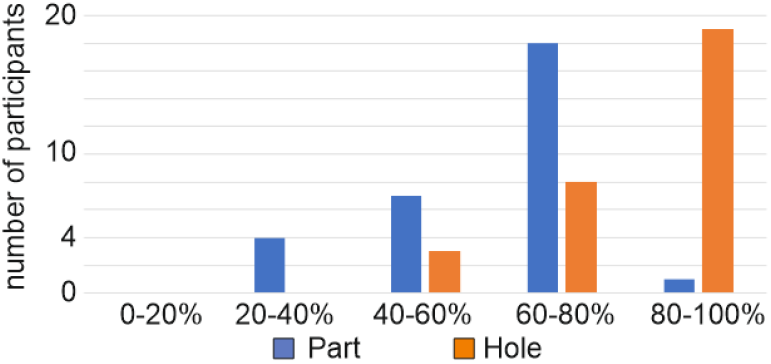
“Close-to” versus “Specific” lookers among participants: The histogram shows the number of participants, divided in ranges of percentage of “Specific” looking at Parts and Holes

Up to this point the question remains what do people actually look at when they perform a so-called “Specific” gaze, or in other words: Is this term justified and why? To address this question we have analyzed all 37680 “Specific” gazes asking at which markers do people look? The result (Figure 8a) plots the averages over all participants and shows a clear over-representation of markers 7,8,9 complemented by slightly higher values for markers 2,3,4 and for 12,13, too. We note that 7,8,9 form the concave corner of the Part/Hole whereby 2,3,4 are the inner-markers of this corner. Hence, it seems that a large fraction of “Specific” gazes is directed at the concave corner. To assess this in more detail, we analyzed potential features consisting of marker combinations. This is motivated by neural processing, because it is known since many years from physiology that line segments as well as corners lead to strong responses of (different) cells in the visual cortex (Hubel and Wiesel 1962). For example, combinations 5,6 or 10,11,12 represent straight line segments, whereas 7,8,9 or 14,5,6 are examples of corners. Panel B of Figure 8 shows that the concave corner and its belonging line-segment are indeed overrepresented by the number of “Specific” gazes. Interestingly none of the outer (convex) corners shows the same. This pattern is the same for Part and Hole. We had also included the inner-markers 1,2,3,4 into this analysis, but there is no clear difference and – to save space – we omit this diagram here.

**Figure 8.**
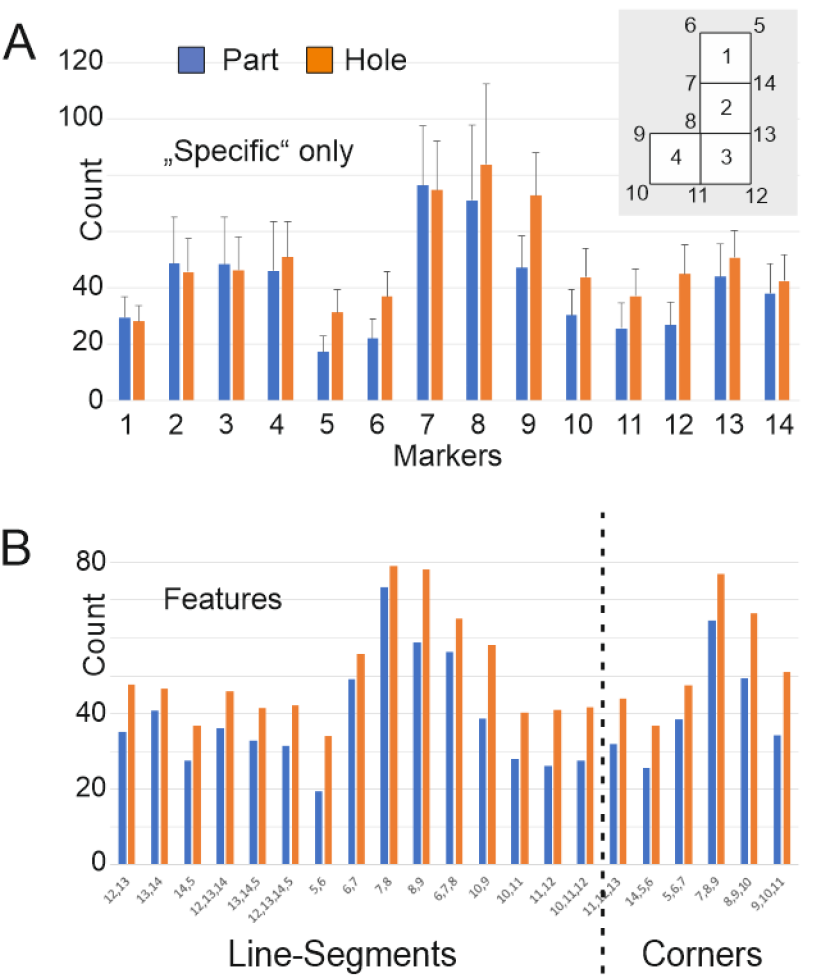
Average values of specific gazes across all participa3n5t7s, A) showing an overview of all marker distributions with higher bars at marker at 7,8, and 9; error bars show confidence intervals of the mean for 30 participants. B) shows this trend on line segments and the corners of the Part/Hole and it also reveals a noticeable over-representation of markers 7, 8, a indicting a large fraction of specific gazes concentrated at the inner corner of the Part/Hole configuration.

### 3.4 Part and hole orientation-based analysis

An important question that arises is whether the observed prevalence of gazes at the concave corner depends on the specific situation. To investigate this, we analyzed all gazes within the “Not middle” subgroup and examined all combinations of Parts and Holes, calculating the quotient of “Specific” gazes at markers 7, 8, and 9 relative to all “Not middle”’ gazes in each scenario, which means: For all given combinations of Part and Hole (e.g., Part=L0 and Hole=L0 in Table 3A, which is on the top left) we calculated how many times participants looked at markers 7,8,9 of the Part divided by how many times “Not middle” gazes at the part happened in general, i.e., at any marker of the considered Part (e.g. L0). Table 3A presents these fractions. For instance, the L0-L0 entry in the top-left corner, 0.13, indicates that for an L0 Part and an L0 Hole, participants looked “Specific” at markers 7, 8, or 9 in 13% of cases and in 87% they looked elsewhere at the Part. The bottom table, in contrast, analyzes gaze patterns at the Holes when different Parts are present. As expected, all values remain below 1, since “Specific-7,8,9” gazes constitute a subset of “All-marker” gazes.

**Table 3.** Quotient of “Specific” gazes at markers 7, 8, and 9, relative to the total “Not middle” gazes for each situation. A) Gaze fractions for Parts in the presence of different Holes, B) gaze fractions for Holes in the presence of different Parts. Yellow color indicates columns (in A) and rows (in B) with lower than the grand-average values. Green color indicates higher than the grand-average values. Asterisks mark significant difference from the grand-average.

|  |  | Looking at Parts (blue) |  |  |  |  |  |  |  |  |  |  |
| --- | --- | --- | --- | --- | --- | --- | --- | --- | --- | --- | --- | --- |
| A |  | L0 | L90 | L180 | L270 | LR0 | LR90 | LR180 | LR270 | AVG | STD |  |
|  | L0 | 0.13 | 0.17 | 0.23 | 0.29 | 0.17 | 0.26 | 0.19 | 0.13 | 0.20 | 0.06 | o |
|  | L90 | 0.11 | 0.16 | 0.26 | 0.33 | 0.16 | 0.33 | 0.19 | 0.08 | 0.20 | 0.09 | o |
|  | L180 | 0.14 | 0.23 | 0.30 | 0.45 | 0.11 | 0.27 | 0.27 | 0.15 | 0.24 | 0.11 | o |
|  | L270 | 0.14 | 0.14 | 0.26 | 0.34 | 0.17 | 0.27 | 0.18 | 0.12 | 0.20 | 0.08 | o |
|  | LR0 | 0.09 | 0.18 | 0.47 | 0.24 | 0.16 | 0.29 | 0.39 | 0.09 | 0.24 | 0.14 | o |
|  | LR90 | 0.16 | 0.20 | 0.26 | 0.36 | 0.12 | 0.24 | 0.20 | 0.13 | 0.21 | 0.08 | o |
|  | LR180 | 0.15 | 0.23 | 0.45 | 0.30 | 0.20 | 0.31 | 0.27 | 0.11 | 0.25 | 0.11 | o |
|  | LR270 | 0.12 | 0.25 | 0.28 | 0.29 | 0.16 | 0.26 | 0.12 | 0.11 | 0.20 | 0.08 | o |
|  | AVG | 0.13 | 0.19 | 0.31 | 0.33 | 0.16 | 0.28 | 0.23 | 0.11 | 0.22 | 0.09 |  |
|  | STD | 0.02 | 0.04 | 0.09 | 0.06 | 0.03 | 0.03 | 0.08 | 0.02 |  |  |  |
|  |  | ** | o | ** | ** | * | * | o | ** |  |  |  |

|  |  | L0 | L90 | L180 | L270 | LR0 | LR90 | LR180 | LR270 | AVG | STD |  |
| --- | --- | --- | --- | --- | --- | --- | --- | --- | --- | --- | --- | --- |
| B | L0 | 0.33 | 0.33 | 0.35 | 0.34 | 0.28 | 0.31 | 0.29 | 0.41 | 0.33 | 0.04 | * |
|  | L90 | 0.38 | 0.30 | 0.32 | 0.36 | 0.32 | 0.31 | 0.39 | 0.34 | 0.34 | 0.03 | * |
|  | L180 | 0.22 | 0.22 | 0.23 | 0.29 | 0.25 | 0.26 | 0.24 | 0.21 | 0.24 | 0.03 | o |
|  | L270 | 0.16 | 0.21 | 0.19 | 0.22 | 0.17 | 0.22 | 0.22 | 0.23 | 0.20 | 0.03 | ** |
|  | LR0 | 0.43 | 0.38 | 0.37 | 0.37 | 0.44 | 0.42 | 0.34 | 0.39 | 0.39 | 0.03 | ** |
|  | LR90 | 0.26 | 0.23 | 0.23 | 0.27 | 0.21 | 0.22 | 0.29 | 0.26 | 0.25 | 0.03 | o |
|  | LR180 | 0.18 | 0.19 | 0.26 | 0.19 | 0.22 | 0.26 | 0.20 | 0.17 | 0.21 | 0.03 | ** |
|  | LR270 | 0.33 | 0.26 | 0.35 | 0.31 | 0.27 | 0.31 | 0.32 | 0.31 | 0.31 | 0.03 | o |
|  | AVG | 0.29 | 0.26 | 0.29 | 0.29 | 0.27 | 0.29 | 0.29 | 0.29 | 0.28 | 0.07 |  |
|  | STD | 0.10 | 0.07 | 0.07 | 0.06 | 0.08 | 0.07 | 0.06 | 0.09 |  |  |  |
|  |  | o | o | o | o | o | o | o | o |  |  |  |

Looking at Holes (blue)

First, we observe that, when analyzing gaze patterns for Parts (Table 3A), the presence of different Holes has little impact, as the fractions within each column remain highly similar (vertical standard deviation ≤ 0.06, with two exceptions). In contrast, for a given Hole, the gaze patterns across different Parts vary more substantially (horizontal standard deviation ≥ 0.06). This indicates that when looking at Parts, the Part itself is the primary factor, rather than the Hole beneath it. Similarly, for Holes (B), the same principle applies: when looking at Holes the gaze behavior is influenced more by the Hole itself than by the Part above it, as evidenced by the smaller horizontal standard deviations compared to the vertical ones.

As previously observed, reaction times were shortest along the equal-piece diagonal—where the Part and Hole share the same configuration (from top left to bottom right)—since these situations are arguably the easiest. Based on this assumption, one might expect fewer “Specific” gazes in these cases compared to others. However, this expectation is not supported by the data, as results along the diagonals exhibit considerable variation.

However, on average, we do observe more “Specific-7,8,9” gazes directed at Holes (B, average 0.28, yellow) compared to Parts (A, average 0.22, yellow). This difference is statistically significant (unpaired t-test, p < 0.001).

Furthermore, we examined whether certain columns or rows deviated significantly from their grand averages. In (A), none of the rows showed such deviations, and similarly, in (B), none of the columns did (see ‘o’ markers on the side or at the bottom). However, several columns in (A) and rows in (B) stood out. Cases with averages significantly higher than their corresponding grand averages are marked in green, indicating a greater-than-average number of ‘Specific-7,8,9’ gazes. Dark colors indicate p < 0.01, while light colors denote p < 0.05.

In Figure 9 we plot histograms of the counts of all specific gazes via columns (for parts) and rows (for holes). Therein we show the gazes corresponding to concave corner in light green color. We mark in Figure 9 with a green arrow (↑) the cases where “Specific-7,8,9” gazes make a substantial part of all specific gazes. Conversely, cases where “Specific-7,8,9” gazes were significantly lower than average are marked with an orange arrow (↓) in Figure 9. Notably, many of the green-arrow cases — where “Specific-7,8,9” gazes were more frequent — correspond to situations where the concave corner of the Parts points downward, whereas for Holes, it points upward.

**Figure 9.**
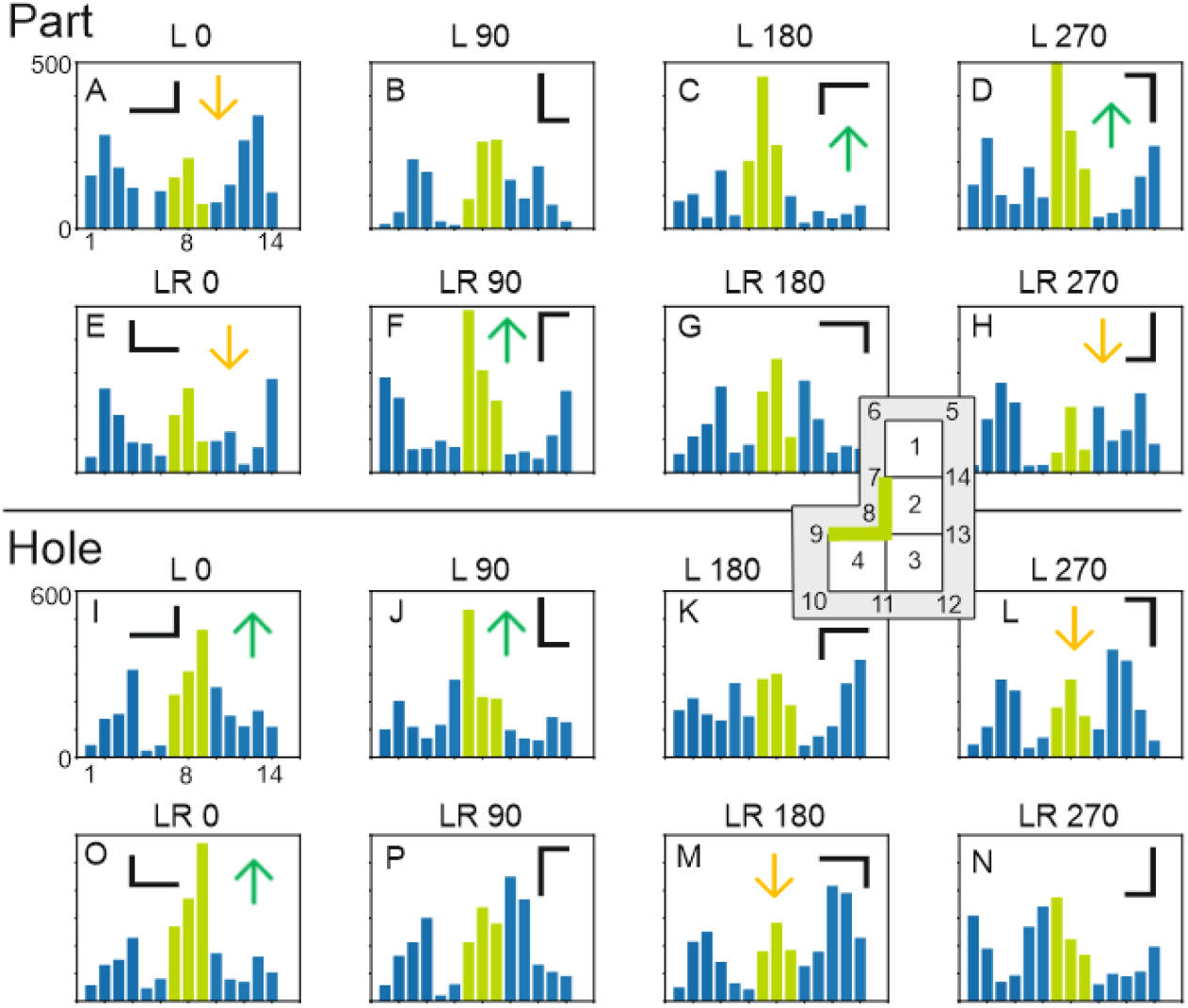
This shows the distribution of gazes at the different markers 1-14 (see inset) where we highlight in light green bars markers in the inset 7,8,9, which are the ones at the concave corner. Arrows depict those cases from Table 3 where significantly more (less) gazes are at the concave corner in green (orange). This figure essentially confirms the results from the table. Sometimes smaller peaks exist at/near markers 1,2,3,4, which are the inner points of the object, but these peaks do not reach any significance level

To quantify this effect, we compared the average values for downward-pointing concave corners versus upward-pointing corners (Table 4) and found a highly significant difference (p < 0.001 in every case). Interestingly, the pattern observed for Holes is the inverse of that for Parts: downward-pointing concave corners in Parts, and upward-pointing concave corners in Holes, elicit more “Specific” gazes than their counterparts.

**Table 4.** Analysis of the proportion of “Specific-7,8,9” gazes for upwards vs downwards pointing concave corners for Parts and Holes.

| **Parts** |  |  |
| --- | --- | --- |
|  | AVG | STD |
| Up | 0.15 | 0.04 |
| Down | 0.29 | 0.08 |
| **Holes** |  |  |
|  | AVG | STD |
| Up | 0.34 | 0.05 |
| Down | 0.22 | 0.03 |

## 4. Discussion

Mental rotation efficiency is influenced by the familiarity and simplicity of geometric forms. Studies indicate that objects with asymmetrical yet predictable structures, such as L-shaped blocks as in our case, are mentally rotated faster and with fewer cognitive demands compared to irregular or complex forms (Tarr and Pinker, 1989), Liesefeld et al. 2015). He et al. (2022) also showed how symmetry in visuospatial structures enhances change detection where participants viewed 3D cube structures and identified changes between symmetrical and asymmetrical designs. Their findings revealed that symmetrical structures improve detection efficiency, though individuals with higher spatial ability performed better overall. These findings also align with Kessler et al. (2018) who showed in a spatial problem-solving study that individuals use stored representations of common shapes to optimize visuospatial processing. This relates to the fact that humans rely on familiar spatial patterns and representations to facilitate problem-solving, recognition, and memory processes (Langlois et al. 2021). All these studies indicate that simpler, or more well-known, geometrical problems are solved more efficiently (and faster) than others. Accordingly, we had hypothesized that participants will recognize a fit between Part and Hole faster and with less errors as compared to a non-fit, where simple configurations will lead to shorter reaction times than more complex ones. Not too unexpectedly, the findings on judgement latency of this study support this conjecture and align, thus, well with the above discussed literature. However more errors in our study were made when judging fit as compared to non-fit.

The detailed analysis of the participants’ gaze pattern, on the other hand, led to interesting findings, too. Table 2 had shown that a fraction of 37% of all gazes landed just in the middle of the canvas, likely indicating the use of peripheral vision to solve the given task. Given the large size of the objects, this is quite conceivable. In other studies, it was also found that adults manage to recognize simple geometric figures using extrafoveal analysis (Chumachenko et al, 2024). Also, fixations around the center of gravity are also known from other tasks, like face and other object recognition (Bindemann et al, 2009, Xu et al, 2014). Alternatively, it was found that people may fixate at blank locations when solving geometrical tasks, which was hypothesized to reduce distractions when thinking about possible solutions (Schindler and Lilienthal, 2019). As our task required much less depth in reasoning as compared to the task of “making a geometric proof” analyzed in the aforementioned study, extrafoveal processing is more likely for our case and is a point for future research.

Of the remaining 63% of gazes there was 1/4^th^ where participants looked “just close enough” at Part or Hole (Table 2, last column). This may indicate a “thrifty” gaze-strategy reducing saccade lengths when looking back and forth at Part and Hole, by not directly looking at them but just close enough. This finding cannot be attributed to any (calibration) errors in the setup as the interspersed calibration controls, where participant had been asked to fixate a certain spot, are highly accurate. Furthermore, we observed that the remaining 3/4^th^ of their gazes was directly targeting the objects at quite specific and discriminable locations. Our findings suggest that the concave, but not the convex, corner of the L plays a special role. To discover this in a rigorous way it had been important to not confound specific looks at the corner markers 7,8,9 with those that are “just close enough” at Part or Hole. Sorting the markers for every situation according to their vicinity to the CoM was addressing this issue. Although some “specific” cases of gazes at 7,8,9 were screened out in this way, at least this procedure ensured that the finding of the prevalence of the concave corner was not just an artifact of chance proximity. In conjunction with this, in an older study we had found that humans divide objects into their parts predominantly at concave junctions (Tamosiunaite et al. 2015) and – even more pronounced – they associate “object-ness” more strongly to those geometrical structures that have only few concavities as compared to those with many (Wörgötter et al. 2015). These findings align with a large set of studies that support the notion that concave structures play a special Gestalt-like role in human object cognition (Koenderink and Van Doorn 1982, Hoffman and Richards 1984, Biederman 1987, Cate and Behrmann 2010, Bertamini and Wagemans, 2013) but also for computer vision (Stein, et al. 2014). Furthermore, we found that also the orientation of the concave corner plays a role for the different gaze patterns. downward-pointing concave corners in Parts and upward-pointing concave corners in Holes, elicit more “Specific” gazes than their counterparts.

## 5. Conclusions

This study contributes on visual perception and object recognition, offering insights into how people interact with their environment through visual cues and shapes. In conclusion, our results shine a light on the actual complexity of the underlying recognition processes. It appears that at least three strategies exist and are used with different prevalence by different people. 1) In many cases (n=30636) this task, which consists of rather large objects, can be solved using just peripheral vision. 2) In a few more cases “Specific” gazes happened (N=37680) focusing mostly of the Gestalt of a concave corner, and 3) less often but still quite pronounced (N=13645) there was the strategy to just look close enough to Part or Hole this way potentially reducing saccadic gaze effort.

Thus, these findings emphasize the importance of proximity, spatial orientation and visual cues in object recognition tasks, indicating that the perceptual strategy employed relies on quite different aspects in the configurations. The measured reactions times confirm the old notion of “simpler is faster”, but it also seems that familiar shapes (e.g., real L) tempt people to react faster but at the same time more often wrongly. The fact that the concave corner attracts gazes is well in hand with several older studies (Koenderink and Van Doorn 1982, Hoffman and Richards 1984, Biederman 1987, Cate and Behrmann 2010, Bertamini and Wagemans, 2013, Tamosiunaite et al. 2015, Wörgötter et al. 2015) but, in addition to this, it appears that participants often operate in an “economic manner” employing peripheral vision instead. Complex vision decision tasks in real life may also be guided by such strategies and future research could expand on these findings by exploring these principles in different real-world applications for improved and safer (error free) object recognition.

## 6. Acknowledgments

This publication was funded by the Deutsche Forschungsgemeinschaft (DFG, German Research Foundation) - Project-ID 454648639 - SFB 1528, “Cognition of Interaction”, Project B01 and by Lower Saxony Ministerium für Wissenschaft und Kultur (MWK), Project: Kognitiv und Emphatisch Intelligente Kollaborierende Roboter (KEIKO), TP6.

## APPENDIX Analysis details

### Camera calibration

The conventional approach to camera calibration involves capturing multiple images of a reference checkerboard pattern. For our purposes this is not feasible and, instead, we used the Tetris grid as a reference pattern. By accurately measuring the pixel positions of the Tetris bricks in several images manually, we then applied the “calibrateCamera” function from OpenCV. The resulting calibration renders a well-defined and undistorted area in the central region of the screen, where the canvas was typically located.

Post-calibration, we exclusively worked with undistorted images to ensure the accuracy of the data we retrieved. The camera matrix was used to map the eye-tracking data onto the undistorted images. By employing the OpenCV “undistortPoints” function from OpenCV, we were able to align the data with the undistorted image space to assure precision of our measurements.

### Processing

To this end, we established a coordinate system with the top left corner of the screen as the origin. The goal was to locate pixel points on the screen and calculate a transformation matrix that effectively projects the pixel coordinates of the video to their corresponding actual locations. For accurate positioning, we utilized four Aruco markers—one situated in each corner of the screen. These markers provide a reliable means of determining orientation and spatial mapping.

Our method started with the identification of the bottom grid in the image. By calculating a preliminary transformation matrix, we established an initial estimate. This matrix enabled us then to direct our search for the exact position of the Part in the video feed. This process was iteratived and this way, we obtained a more accurate transformation matrix, utilizing pixel data from both the bottom grid and the input Part. Each transformation matrix was recorded in a JSON file for future reference.

Next we needed to detect the black boxes that constituted the bottom grid. We implemented a systematic series of steps: first, we converted the image to grayscale. Then, we focused on the specific pixel range of interest—dark pixels between 0 and 50—amplifying them in our analysis. We applied edge detection algorithms to create clear boundaries around the boxes followed by a dilation process to enlarge these edges, ensuring robustness against potential gaps in detection.

Once the boxes were delineated, we quantified the pixels within each detected area, identifying those measuring between 500 and 1800 pixels as viable candidates. Following this, we computed the median distance between these locations to estimate the grid spacing, filtering out locations lacking nearby neighbors within this distance. With this analysis we extracted the largest connected cluster, which represented the grid layout.

The positioning and orientation of the input Part is straightforward, achieved by analyzing the pixel density across two regions on the screen’s upper half. By comparing the number of dark pixels on each side, the system can quickly identify the location of the Part.

However, for the Hole location, the procedure necessitates a careful scan of the grid below the input area. By examining each potential grid location for a match in piece geometry, the system effectively narrows down possibilities. Central to this process are lookup tables established for all Holes. Each shape is represented through a series of coordinate points relative to a designated center, allowing for transformations that include mirroring and rotation. The coordinates for the L-block in Figure 5 illustrate this method effectively: each point is annotated by a unique numeric identifier, ranging from 1 to 14. By utilizing a function that transforms these block numbers into screen coordinates, we ensure that each figure is accurately represented on the display, regardless of its orientation.

Once the transformation is complete, resizing and translating these coordinates into screen positions are stored in a JSON file, a structured format ideal for subsequent processing. This file serves as a repository of all critical positional data, thereby facilitating smooth integration into further analyses of our visual tasks.

## Footnotes

1 A tetromino is a geometrical figure consisting of 4 squares, which are attached to each other with their sides touching, forming, for example, an „L” or any other possible figure .

